# Extraction of biological signals by factorization enables the reliable analysis of single-cell transcriptomics

**DOI:** 10.1101/2023.03.04.531126

**Authors:** Feng Zeng, Xuwen Kong, Fan Yang, Ting Chen, Jiahuai Han

**Affiliations:** Department of Automation, Xiamen University, Xiamen, Fujian 361102, China; State Key Laboratory of Cellular Stress Biology, School of Life Sciences, Faculty of Medicine and Life Sciences, Xiamen University, Xiamen 361102, China; Research Unit of Cellular Stress of CAMS, Cancer Research Center of Xiamen University, School of Medicine, Faculty of Medicine and Life Sciences, Xiamen University, Xiamen, Fujian 361102, China; National Institute for Data Science in Health and Medicine, Xiamen University, Xiamen 361005, China; Institute for Artificial Intelligence, Department of Computer Science and Technology, Tsinghua University, Beijing 100084, China

## Abstract

Accurately and reliably capturing actual biological signals from single-cell transcriptomics is vital for achieving legitimate scientific results, which is unfortunately hindered by the presence of various kinds of unwanted variations. Here we described a deep auto-regressive factor model known as scPheno^XMBD^, demonstrated that each gene’s expression can be split into discrete components that represent biological signals and unwanted variations, which effectively mitigated the effects of unwanted variations in the data of single-cell sequencing. Using scPheno^XMBD^, we evaluated various factors affecting IFN *β* -stimulated immune cells and demonstrated that biological signal extraction facilitates the identification of IFN*β*-responsive pathways and genes. Numerous experiments were conducted to show that scPheno^XMBD^ could be utilized successfully in enhancing cell clustering stability, obtaining identical cell populations from diverse data sources, advancing the single-cell CRISPR screening of functional elements, and minimizing the influence of inter-subject discrepancies in the cell-disease relationships. scPheno^XMBD^ is anticipated to be a dependable and repeatable method for the precise analysis of single-cell data.

---

The advancement of single-cell RNA sequencing (scRNA-seq) during the past decade has substantially improved our understanding of the cellular and molecular features at individual cell level^1–3^. However, without any exception all scRNA-seq data contain unwanted variations, the variations that are not related to task of a given study. Unwanted variations can be caused by many reasons, some of them can systematically distort gene expression measurements, including cell cycle, sequencing depth, gene capture rate, batch effects^4,5^, and inter-subject variations^6,7^. Consistence and accuracy are concerns for the fundamental analysis tasks like cell clustering and data integration when unwanted variations present^8,9^. In fact, unwanted variations can lead to false identification of previously undiscovered cell types; unwanted variations can also reduce the ability to recognize cell populations associated with specific phenotypic abnormalities. It is well known that the accuracy and replicability of the results deduced from scRNA-seq heavily rely on the removal of unwanted variations, but current tools cannot do it satisfactorily.

It is possible to adjust bulk transcriptomics basing on internal references like housekeeping genes or molecular spike-ins that are added in fixed amounts^10^. But these cannot be applied to scRNA-seq as each gene’s expression can vary considerably from cell to cell. Instead, Seurat^11^ and other data-processing algorithms^12^ typically integrate normalization^13^, dimensional reduction^14^, and batch correction^15,16^ to eliminate unwanted variations among cells and adjust for batch effects. Given the impossibility to predict the source and nature of all unwanted variations, these data-processing techniques suffer from over-correction and under-correction frequently^17,18^. The former could misinterpret vital biological signals as unwanted variations, causing them to be disregarded. The latter will preserve a portion of unwanted variations as spurious biological signals. Downstream analyses can be damped by both.

While desired variations in scRNA-seq data are those that are linked to biological phenotypes under study, these wanted variations in gene expression may be inferred differently across distinct phenotypes, levels of specificity, or scales, depending on the method of calculation employed. Therefore, it is important to carefully consider the computational methods used in analyzing scRNA-seq data to ensure that the inferred variations accurately reflect the biological phenomenon under investigation. For example, clustering with coarser parameters suggests gene expression variations on a smaller scale, whereas clustering with finer parameters shows gene expression variations on a wider scale. By integrating diverse phenotypes with scRNA-seq data, it is possible to build a precise representation of biological signals, which could enhance precision and replicability of scRNA-seq data analyses. To date, methods for this purpose are absent.

We here introduce scPheno^XMBD^, a deep auto-regressive factor model that is used to extract the biological signals imbedded in transcriptome, identify gene expression variations associated with each of the phenotypes, and re-build the accumulative effect of multiple phenotypes on cell states. scPheno^XMBD^ is adept at eliminating unwanted variations in gene expression and reconstructing the noise-free transcriptome, hence preserving the integrity of the original biological signals. The evaluation of scPheno^XMBD^ demonstrated that extracting biological signals is advantageous in that it provides a more precise understanding of the pathways’ responses, a dependable cell clustering and data integration, a mitigation of undesired effects of confounding factors such as cell cycle and replicates, a reduction of discrepancies among individual samples. scPheno^XMBD^ should be applicable across a broad range of fields.

## Results

### Overview of scPheno^XMBD^

In general, scPheno^XMBD^ repeatedly engages in a self-teaching game of *deconstruction-and-reconstruction* to learn how to factorize gene expression into distinct parts that resulted from different biological signals and unwanted variations (Supplementary Fig. 1). The deconstruction of gene expression is accomplished by using an array of factorization neural networks. A factorization neural network will factorize gene expression pertaining to a phenotypic factor and project cells onto a latent variable space, where the latent variable specifies a hidden cell state and cells of the same hidden states (i.e., phenotypic properties) will cluster together (Fig. 1a). We apply the classification and regression criteria to ensure that gene expression relating to phenotypic changes can be accurately retrieved and factored. On the factorized latent variable spaces, an array of multiple-layer neural network classifiers is constructed for phenotypic predictions. If a factorization is valid, the classifier that follows must reliably predict the phenotypic labels of cells. Then, a deep factor model is constructed to regress gene expression against phenotypic factors, such cell type, an intriguing cell phenotype, batch, and others (Fig. 1b). Using solely phenotypic input, including phenotypic labels and prediction feedbacks, the deep factor model will infer the factorized latent variable spaces. Following that, a neural network is utilized to rebuild the gene expression of cells based on the combination of factorized latent variables. If single-cell transcriptional profiles are reliably recreated, then gene expression in relation to phenotypic variables can be properly retrieved. Intriguingly, after model training, the factorization neural networks and the reconstruction neural network can be coupled to predict gene expression in relation to any factor combination (Fig. 1c). This will certainly pave the way for deciphering the intricate interactions between multiple factors and gene expression. In sum, the whole system acts like auto-regression that is to use scRNA-seq dataset(s) to predict factorizations and their related phenotypes, followed by using phenotypic annotations and prediction feedback to reconstruct scRNA-seq dataset(s). Consequently, scPheno^XMBD^ is best described as the deep auto-regressive factor model.

**Figure 1.**
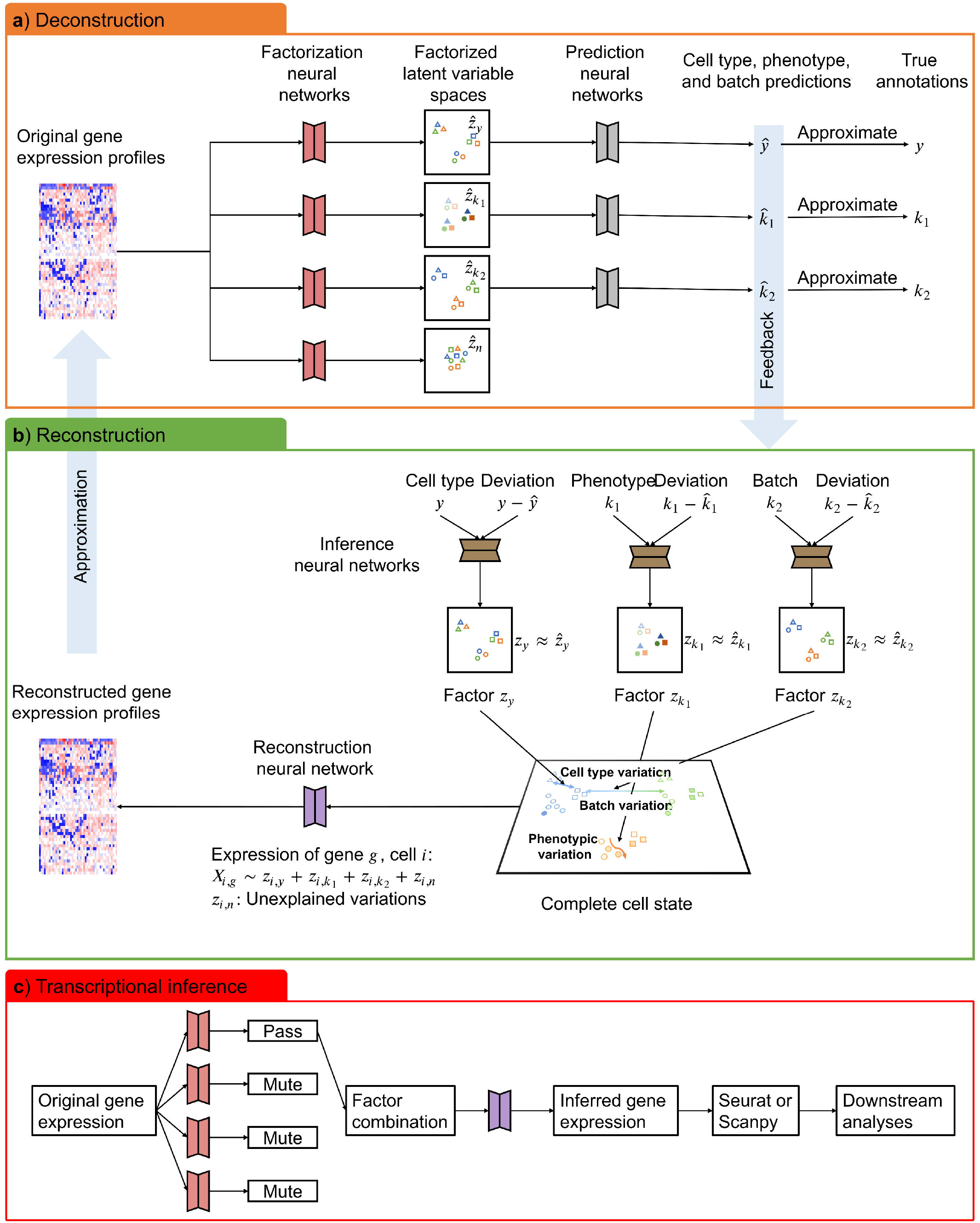
scPheno^XMBD^ reconstructs gene expression of cells by controlling factors. **a**, Given cells’ transcriptional profiles, scPheno^XMBD^ decomposes the variations in gene expression associated with distinct factors, each of which corresponds to a latent state space, for example, cell type, cell phenotype, and data batch. Prediction neural networks are constructed to predict the labels of different aspects. **b**, scPheno^XMBD^ reconstructs the state of a cell in each latent state space by inferring the standard cell state (i.e., archetype) from the given annotated label and then inferring the deviation from the archetype based on the difference between the annotated and predicted labels. The latent states pertaining to distinct factors are combined as the complete cell state and then used for the reconstruction of gene expression. **c**, scPheno^XMBD^ enables the reconstruction of gene expression corresponding to the combinatorial effect of various factors, which can then be analyzed with standard methods.

### Gene expression is factorizable

We used scRNA-seq data of interferon beta (IFN *β*) stimulation of human peripheral blood mononuclear cells (PBMCs)^19^ to demonstrate that the expression level of each gene can be factorized into subcomponents, with each subcomponent indicating the effect of a phenotypic factor. This dataset contains 7,451 IFN*β*-stimulated cells and 6,548 control cells (Fig. 2a). The uniform manifold approximation and projection (UMAP)^14^ plot reveals a correlation between the distribution of PBMC cells and three factors, namely cell type, cell subtype, and stimulation status (Fig. 2b). We used scPheno^XMBD^ to assess the effects of these factors (Fig. 2c). The bulk of differences in gene expression between cells are best explained by cell type, followed by cell subtype and stimulation status (Fig. 2d), and the impact of unknown factors is small. This means the majority of changes in this dataset can be factorized into three subcomponents: cell type, cell subtype, and stimulation status.

**Figure 2.**
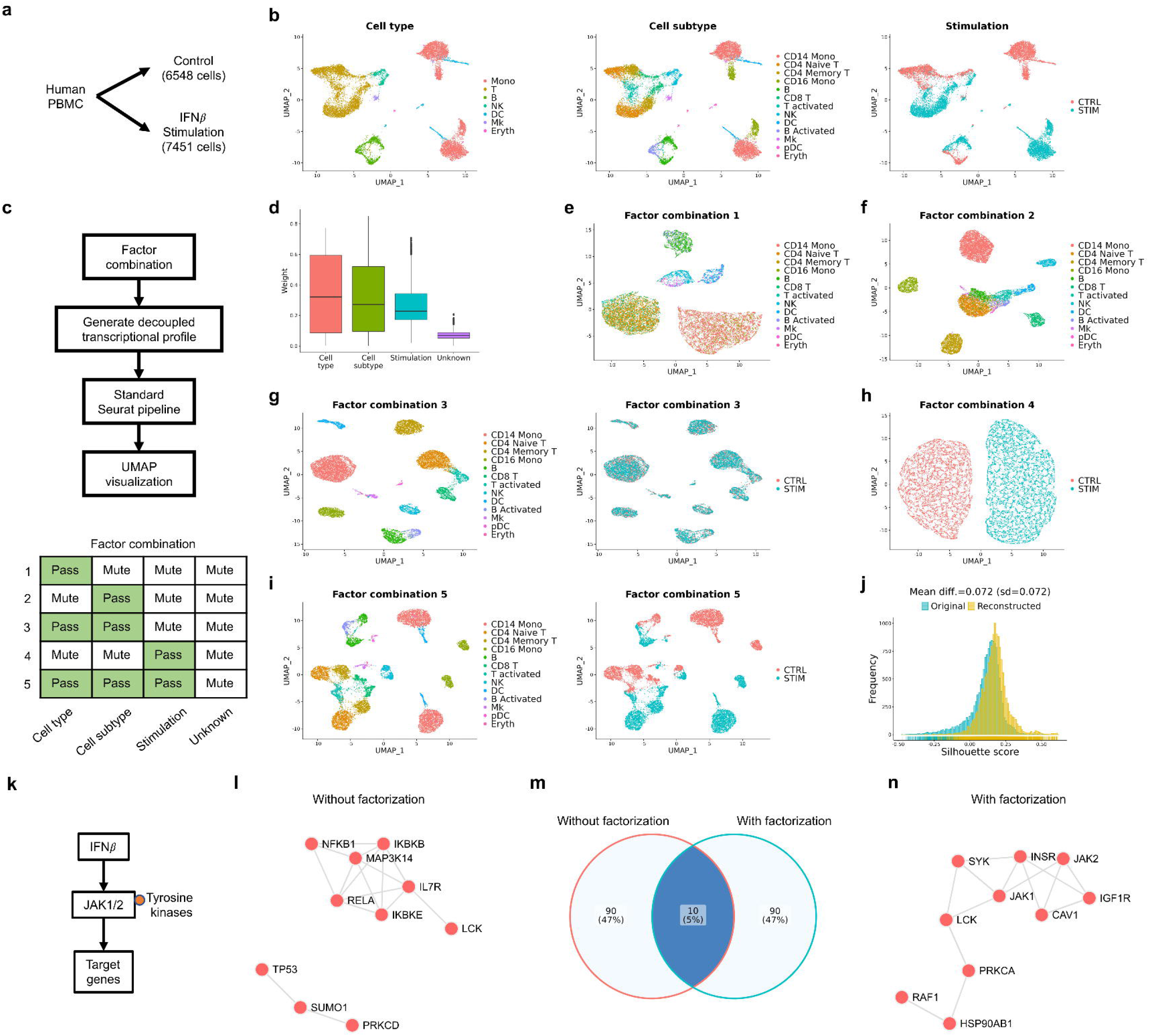
Factorizing scRNA-seq data of IFN*β* stimulation and inferring gene expression profiles corresponding to different combinations of factors. **a**, A scRNA-seq data of 6,548 control cells and 7,451 IFN*β*-stimulated cells from a human PBMC research^19^ were utilized. **b**, The characteristics of the original scRNA-seq data with respect to three factors, namely cell type (left), cell subtype (middle), and stimulation status (right). **c**, Schematic of the experiments for the validation of scPheno^XMBD^. **d**, The estimated weights of different factors contributing to the variability of the scRNA-seq data. **e-g**, The UMAP visualization of cell distributions derived from the reconstructed gene expression for cell type (**e**), cell subtype (**f**), and the combination of the two (**g**). **h**, The UMAP visualization of cell distributions based on the reconstructed gene expression associated with stimulation status. **i**, The reconstructed data with respect to the combinatorial effect of cell type, cell subtype, and stimulation status preserves the property of the original data. **j**, Compare the structures of cell populations revealed by the original and reconstructed data separately by using cells’ Silhouette scores. The average difference between cells’ Silhouette scores is 0.072±0.072. **k**, Illustrating the signaling pathway of IFN*β* stimulation. **l**, The overrepresented proteins estimated from the original data by using EnrichR. **m**, Comparison of top 100 differentially expressed genes estimated from the original data and the reconstructed data regarding stimulation status. **n**, The overrepresented proteins estimated from the reconstructed gene expression for stimulation status by using EnrichR.

We therefore used scPheno^XMBD^ to factorize the IFN*β*-stimulated PBMC dataset and then reconstructed transcriptsome dataset by factor combinations. Seurat^20^ was fed on the reconstructed dataset for routine analysis such as the visualization of gene expression factorization using UMAP (Fig. 2c). Evaluations of three-factor combinations were conducted. First, we factorized and retrieved independently the transcriptional signals associated with cell type and cell subtype. Using the transcriptional signals pertaining to cell type, the UMAP plot showed that cells of distinct cell types are segregated while cells of different subtypes are intermingled within a cell type (Fig. 2e). Using the transcriptional signals pertaining to cell subtype, the UMAP plot revealed that cells are dispersed according to cell subtypes rather than cell types (Fig. 2f). The distribution of cells precisely reflected the relationships between cell types and cell subtypes when both factors were examined (Fig. 2g). Second, we removed the transcriptional signals pertaining to cell type and subtype and focused on the effect of stimulation status alone. Cells were divided into two groups (Fig. 2h). One contained IFN*β*-stimulated cells, while the other contained control cells. Third, we eliminated noise and counted on the interaction between cell type, cell subtype, and stimulation status to reconstruct the transcriptional signals. As shown in Supplementary Fig. 2, no discernible correlation was found between the observed variations among cells in the noise latent space and the three factors considered. Furthermore, the reconstructed dataset demonstrated a high degree of fidelity to the essential properties of the original scRNA-seq dataset, as evidenced by Figs. 2b and i. Notably, the cell population structures between the original and reconstructed data were found to be highly similar, with an average Silhouette score of 0.072±0.072 (Fig. 2j). Thus, the factorizability of gene expression is achievable, and scPheno^XMBD^ provides a straightforward and adaptable means for observing several features imbedded in transcriptomes.

### The signals of a specific biological process are extractible

Next, we confirmed that it is possible to extract signals unique to a biological activity from transcriptomes. The UMAP reconstructed from the transcriptional signals pertaining to cell type and cell subtype revealed that the stimulation labels were mixed (Supplementary Figs. 3 and 4). Similarly, the labels of both cell type and cell subtype were totally intermingled in the UMAP obtained from the stimulation-related transcriptional signals (Supplementary Fig. 5). These results demonstrated that it is feasible to separate transcriptional signals for a particular activity.

We then tested whether the separation of transcriptional signals helps the functional investigation of IFN*β* stimulation. IFN*β* exerts a variety of effects on immune cells, including the regulation of growth and proliferation, the initiation of anti-viral reactions, and the coordination of pro-inflammatory and anti-inflammatory responses. Studies have well established that IFN*β* stimulates JAK-STAT signaling pathway to initate IFN responsible gene expression (Fig. 2k), and the JAK-STAT can also engage NF κ B signaling pathway that contributes to the regulation of cell proliferation and stress response^21^. Due to the intricacy of the biological processes induced by IFN*β*, NFKB1, NFKBIB, RELA, and IKBKE, which are the essential components of NFκB signaling pathway^22^, were overrepresented in the top 100 upregulated genes in the original data (Fig. 2l). In other words, the major antiviral and inflammatory genes were not revealed by directly analyzing the original data. We used scPheno^XMBD^ to separate the combined effect of proliferation (characterized by cell type and cell subtype) and inflammation (characterized by stimulation status). We retrieved gene expression regarding stimulation status while excluding the transcriptional signals associated with cell type and cell subtype. The top 100 genes reported by differential analysis of the extracted gene expression were vastly different from those discovered in the original scRNA-seq dataset (Fig. 2m), which had the proteins of JAK-STAT pathway, such as JAK1, JAK2, and tyrosine proteins (e.g., SYK and LCK) (Fig. 2n). As shown by these analyses, complexity of a system may affect the specificity of functional analysis of transcriptional signals. By identifying transcriptional signals pertaining to an aimed biological process, scPheno^XMBD^ may minimize system complexity and increase the specificity of functional analysis.

### Multi-scale information improves the consistence of cell clustering

Cell clustering is crucial for the investigation of cell heterogeneity. But stochastic unwanted variations prevent cell clustering from producing a consistent estimation of the cell-cell relationships. Even if a clustering method is executed with optimal parameters, utilizing different numbers of genes could generate incongruous results.

scRNA-seq data of human fibroblast cells^23^ was utilized to showcase the approach. Genes were ranked decreasingly according to the expression variance across cells. Then, different numbers of top genes were selected as highly variable genes (HVGs) for cell clustering. It is observed that the relationships between cells were random when few HVGs were used (Fig. 3a, left), whereas the determined relationships between cells were achieved when more HVGs were used (Fig. 3a, right). The adjusted Rand index (ARI) was used to assess the consistence between clustering results. The pairwise comparison revealed significant variations in clustering outcomes when using unprocessed data (Fig. 3b). An array of multi-scale processed datasets was generated. Specifically, the multi-scale approach involved collecting the clustering results of different sets of HVGs as the phenotypes regarding varied degrees of specificity. This informed scPheno^XMBD^ to extract the scale-dependent variations in gene expression (Fig. 3c). The results were inspiring that the multi-scale processed data significantly enhanced the consistence between cell clustering results (Figs. 3d and e). These findings suggest that utilizing the multi-scale processing strategy may help overcome the inconsistences observed in clustering results, particularly when dealing with complex biological datasets.

**Figure 3.**
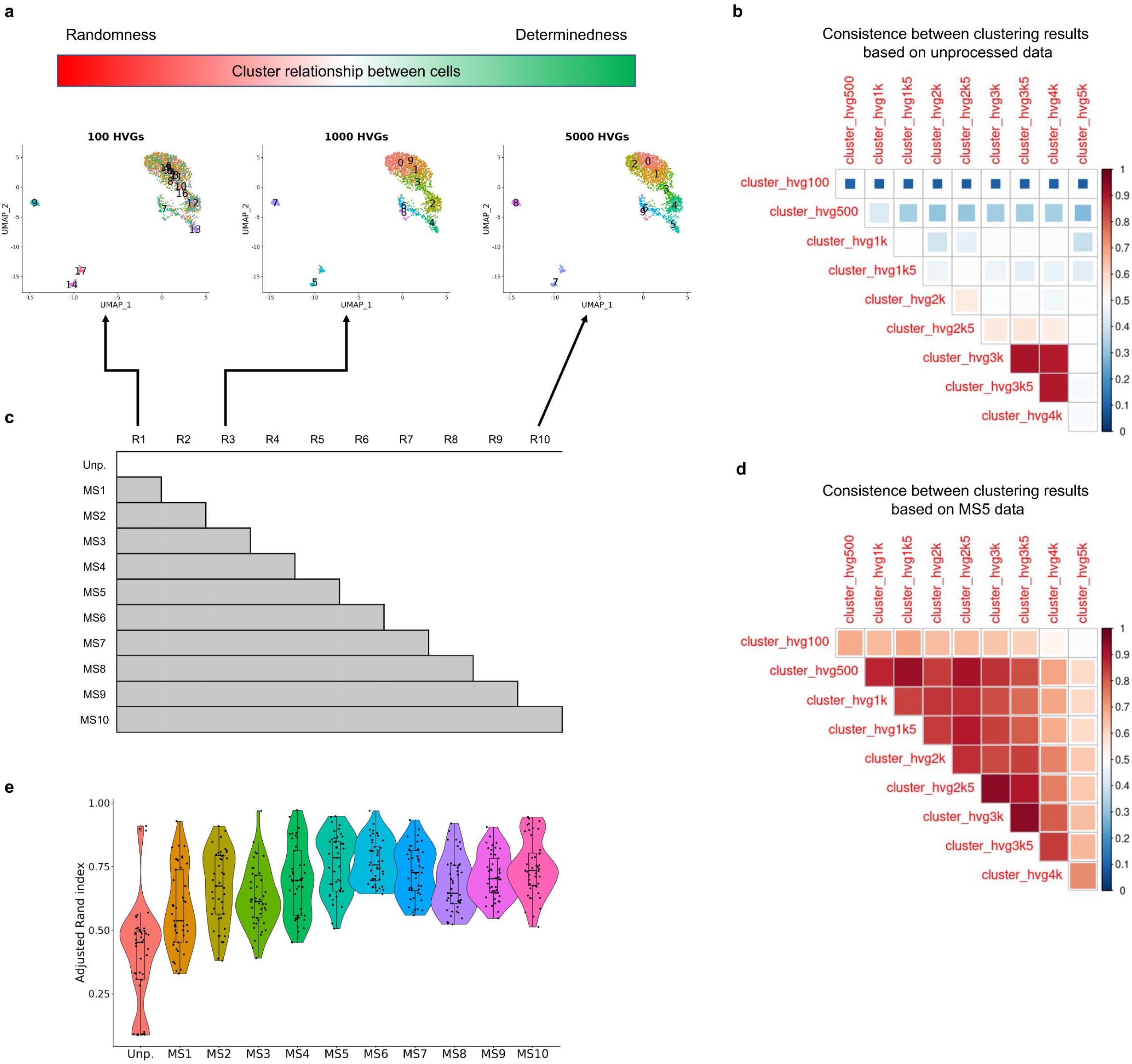
Integrating multi-scale transcriptional variations enhances the consistence of cell clustering. **a**, The results of clustering human fibroblast cells^23^ with varying numbers of HVGs range from 100 to 5,000, with a 500-gene increment. Three results are displayed. **b**, Assessment of the consistence between each paired clustering results estimated from the unprocessed data. **c**, Schematic illustration of the generation of multi-scale processed datasets represented as MS1 to MS10, which integrate different sets of clustering labels. **d**, An example of MS5 exhibits the multi-scale processed data generate more consistent clustering results. **e**, The ARI scores for all multi-scale processed datasets.

### By decomposing wanted and unwanted variations of the data from each of the participants to exclude inter-subject discrepancy

Besides random noises, unwanted variations such as inter-subject discrepancy could systematically distort gene expression readouts and introduce excessive unexpected heterogeneity to cells collected from different individuals, thereby compromising the replicability of scRNA-seq data. For case-control studies, these unwanted variations can severely impede the ability to identify the alterations in cells induced by disease and hinder the precise interpretation of scRNA-seq data. We retrieved CD14+ monocyte cells in the bloodstream of 4 healthy participants and 5 severe influenza patients from the data of a COVID-19 single-cell study^24^ to illustrate the effect of inter-subject variations (Fig. 4a). The differences among each of the participants in either healthy or infected group lead to a substantial inter-individual heterogeneity in CD14+ monocyte cells (Figs. 4b and c). As a result, it was difficult to determine the severe influenza infection caused alterations in CD14+ monocyte cells (Fig. 4d). Most single-cell case-control studies use batch correction algorithms such as canonical correlation analysis (CCA) of Seurat to eliminate the random differences between individuals. However, current batch correction algorithms cannot address the combinatorial effect of multiple factors on cell heterogeneity. Although batch correction algorithms could be used to compensate for subject-to-subject variability (Figs. 4e and f), the differences in cells between health and severe influenza infection could be removed too (Fig. 4g). The over-correction made it challenging to identify disease-associated cells. Here we used scPheno^XMBD^ to decompose the inter-subject and severe influenza infection-related variations in gene expression. After eliminating transcriptional differences between participants (Figs. 4h and i), the influenza-responsive CD14+ monocyte cells were clustered together and distinguished from CD14+ monocyte cells in healthy participants. The identical cell populations were obtained in two conditions, allowing the precise identification of the prevalent alterations by severe influenza infection on patients (Fig. 4j).

**Figure 4.**
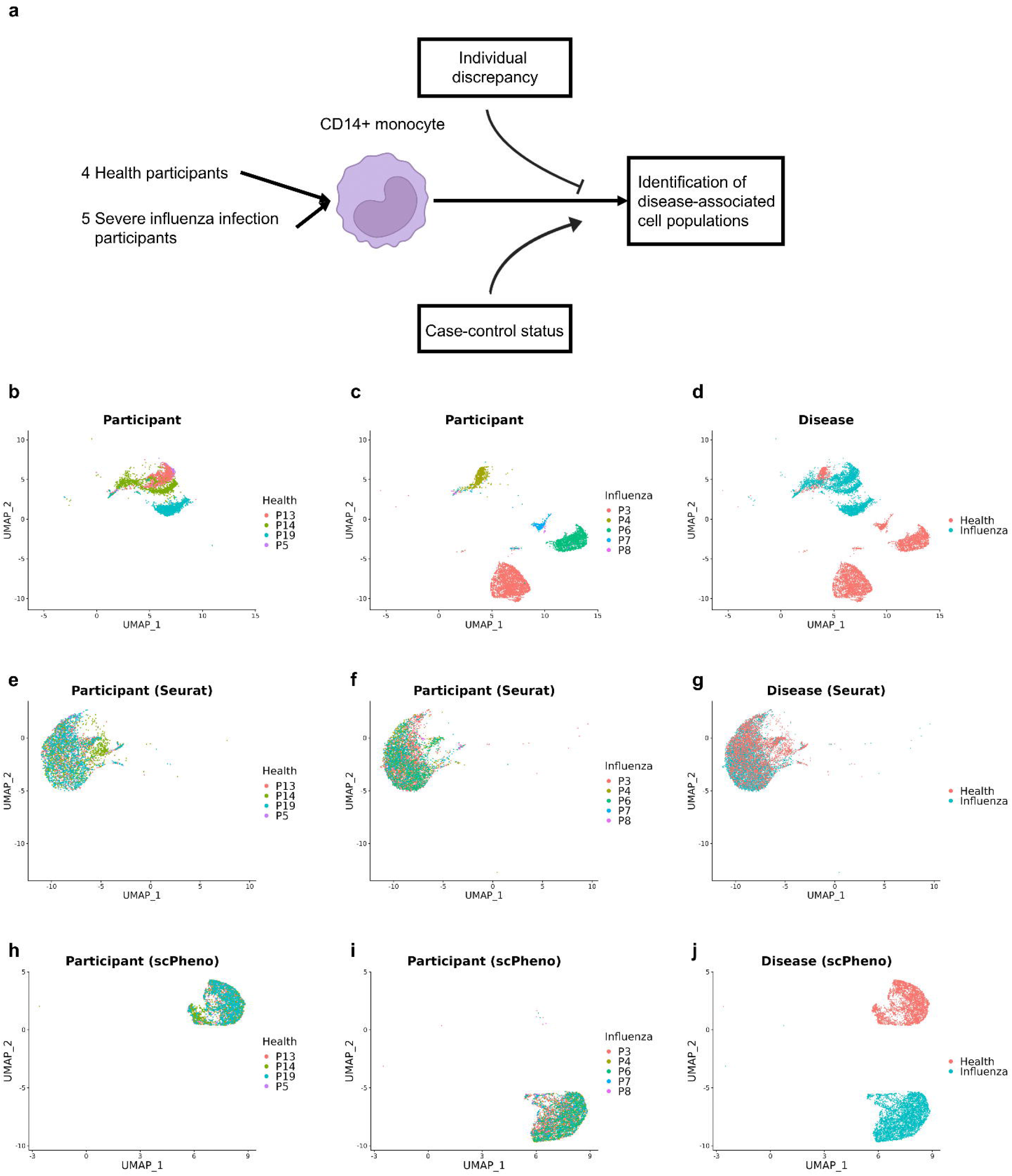
Mitigating the effect of inter-subject variations on the identification of disease-associated cells. **a**, CD14+ monocyte cells from the peripheral blood of 4 health participants and 5 severe influenza patients^24^ were utilized. In case-control studies, inter-subject discrepancy could hinder the characterization of disease-associated cells. **b**,**c**, The impact of the random differences between individuals on CD14+ monocyte cells during health state (**b**) and severe infection state (**c**). **d**, The variability of CD14+ monocyte cells from healthy participants and severe influenza patients. **e-g**, Seurat was used to eliminate the differences between individuals. The distributions of cells for healthy participants (**e**), severe influenza patients (**f**), and disease conditions (**g**). **h**,**i**, scPheno^XMBD^ was used to eliminate the random differences among healthy participants (**h**) and among severe influenza patients (**i**). **j**, scPheno^XMBD^ retained the variability of cells corresponding to illness.

### scPheno^XMBD^ offers the potential for generalizable cellular discovery

Traditional transfer learning methods involve adapting existing models to new datasets based on shared information^25^, which may obscure the natural variations among cells and hinder the development of broad knowledge. However, we postulate that the use of extracted biological signals for cellular discovery may enable generalization to previously unexplored datasets. To validate this generalizability, we employed the scPheno^XMBD^ model, trained using the Seok et al. dataset^24^ of Korean participants, to analyze the Ahern et al. dataset^26^ of 14,689 and 5,330 CD14+ monocyte cells collected from 10 healthy and 12 severe influenza-infected British participants (Fig. 5a). Our results indicated that 82.4% of CD14+ monocyte cells from healthy participants were correctly mapped to the normal CD14+ monocyte cluster identified in the Seok et al. dataset, while approximately 16.2% were erroneously classified as abnormal (Fig. 5b). This categorization was affected by sequencing quality (Fig. 5c), but could be effectively controlled through gene number-based filtration (Fig. 5d). Identical results were obtained for the influenza-responsive CD14+ monocyte cells in the Ahern et al. dataset (Figs. 5e-g). Encouragingly, there were no inter-subject variations in the scPheno^XMBD^ results (Supplementary Fig. 6). This confirms the generalizability of scPheno^XMBD^’s knowledge of how influenza infection affects CD14+ monocyte cells from the Seok et al. dataset. To further verify the generalizability, we also trained a scPheno^XMBD^ model with the Ahern et al. dataset and applied it to the Seok et al. dataset, yielding consistent results (Supplementary Fig. 7).

**Figure 5.**
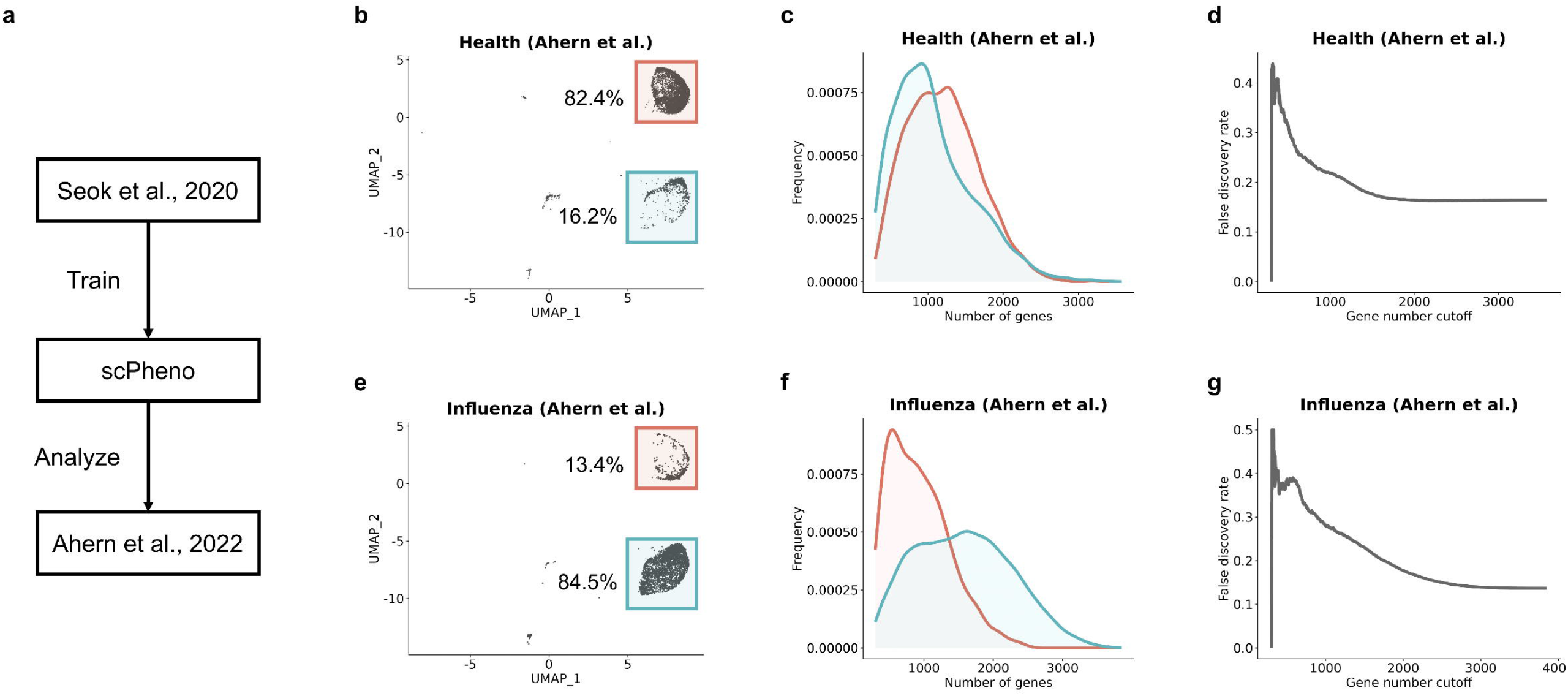
scPheno^XMBD^ is generalizable to unexplored data. **a**, The scPheno^XMBD^ model trained with the Seok et al. dataset^24^ is applied to the Ahern et al. dataset. **b**, The embedding of CD14+ monocyte cells from the healthy participants in the Ahern et al. dataset^26^. The proportions of the correctly classified monocytes and misclassified CD14+ monocyte cells are approximately 82.4% and 16.2%, respectively. **c**, The distribution of gene numbers in the correctly and incorrectly classified CD14+ monocyte cells. **d**, Illustrating false discovery rate varies with gene number-based filtration. **e**, The embedding of CD14+ monocyte cells from the influenza infection patients in the Ahern et al. dataset. The proportions of the correctly classified monocytes and misclassified CD14+ monocyte cells are approximately 84.5% and 13.4%, respectively. **f**, The distribution of gene numbers in the correctly and incorrectly classified CD14+ monocyte cells. **g**, Illustrating false discovery rate varies with gene number-based filtration.

### Dissecting cellular regulation network

Genetic perturbation of genes is one of the most effective approaches to study cellular regulation networks. Unfortunately, gain or loss of function of one gene could have effect on multiple regulatory pathways. Here we use the application of scPheno^XMBD^ in analyzing the data of a single-cell CRISPR screening as an example to test whether this tool can help the study of gene regulatory networks.

Single-cell CRISPR screening is an approach to reveal the overall effect of every gene because single-cell sequencing can read tens of thousands transcriptional signals in a single CRISPR-edited cells (that are perturbed). However, single-cell CRISPR screening is affected by a variety of known and unknown confounding factors. Few computational methods exist for handling confounding factors in CRISPR screening of single cells. We reanalyzed a single-cell CRISPR screening dataset that characterizes regulatory genes in THP1 cells^27^ (Fig. 6a). The analysis of single-cell CRISPR screening dataset was found to be significantly influenced by replicates and cell cycle (Figs. 6b and c). First, the distribution of THP1 cells primarily reflected the influence of these two confounding factors as opposed to the CRISPR perturbation (Fig. 6d). Second, a large fraction of CRISPR-edited cells could not be separated completely from unperturbed cells (Fig. 6d). Since a hierarchy of genes is responsible for modulating the cellular response to IFN *γ*, decitabine, and TGF*β*1 activation, including interferon receptors (e.g., IFNGR1 and IFNGR2) and JAK-STAT signaling mediators (e.g., JAK2 and STAT1)^21^, it is logical that genes at different stages of signal transduction would exert diverse impacts on the cellular response to IFN*γ*. However, THP1 cells devoid of IFNGR1, IFNGR2, JAK2, and STAT1 revealed comparable distributions (Fig. 6e). We used scPheno^XMBD^ to factorize transcriptional signals and exclude unwanted variations pertaining to replicates and cell cycle. The replicates and cell cycle effects appear to have been successfully removed from the processed dataset (Figs. 6f and g). The distinctions between the CRISPR-edited cells and unperturbed cells became apparent (Fig. 6h). In addition, THP1 cells devoid of IFNGR1, IFNGR2, JAK2, and STAT1 were distributed differently (Fig. 6i). The processed dataset could also assist in the identification of novel regulatory genes, including SMAD4, IRF1, STAT2, and BRD4 (Fig. 6i). scPheno^XMBD^ provides a prominent approach for processing single-cell CRISPR screening datasets and will facilitate the application of gene regulatory networks created from CRISPR-edited cells.

**Figure 6.**
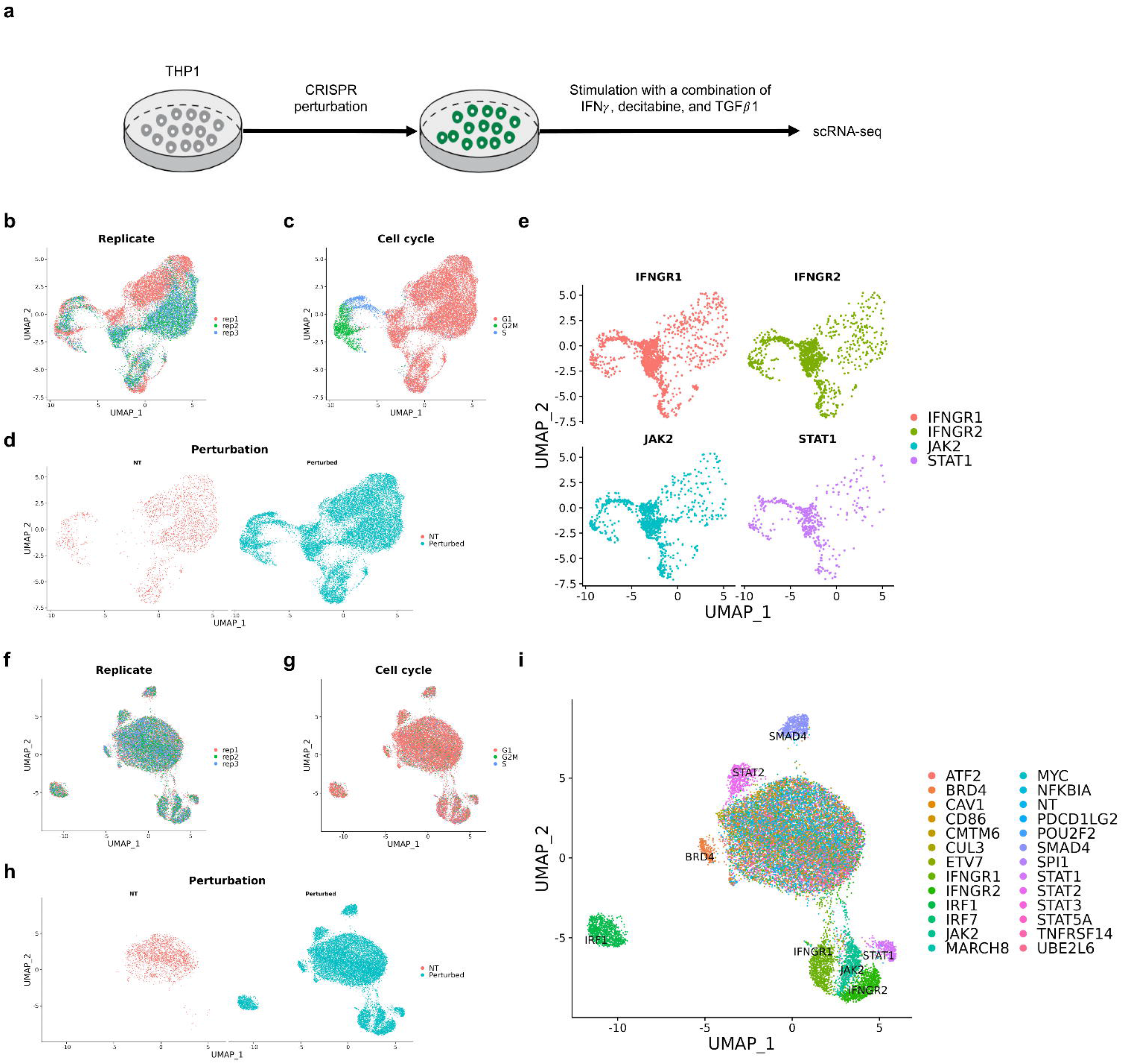
scPheno^XMBD^ improves single-cell CRISPR screening. **a**, Schema illustrating the generation of scRNA-seq data for the CRISPR-edited THP1 cells^27^. **b-d**, Cell distribution correlates with replicate (**b**), cell cycle (**c**), and perturbation (**d**). **e**, Visualization of the comparable distributions of cells devoid of IFNGR1, IFNGR2, JAK2, and STAT1. **f-h**, Illustration of the effects of replicate (**f**), cell cycle (**g**), and perturbation (**h**) estimated from the scPheno^XMBD^-reconstructed data. **i**, scPheno^XMBD^ reveals that the CRISPR-edited cells with different impacts distribute at separate locations.

## Discussion

In single-cell transcriptomics, unwanted variations and biological signals co-exist, raising the replicability and reliability crises. Extensive attempts have been made to eliminate as many unwanted variations as feasible. However, the origins and characteristics of unwanted variations vary from study to study, and some unwanted variations remain unexplored. Hence the elimination of unwanted variations has limited effectiveness and practice application. The extraction of biological signals is an alternate method for overcoming the replicability and reliability crises, but no such approach has been reported. Using factor modeling and deep probabilistic modeling, here we developed a theoretical and methodological framework for the factorization of biological signals and unwanted variations in single-cell transcriptomics. The advancements made by our efforts are listed below. First, our work demonstrate that gene expression is multidimensional and can be factored into subcomponents that correspond to biological signals and unwanted variations. Second, transcriptional signals of a particular biological process can be extracted. Third, the extraction and reconstruction of datasets according to biological signals greatly improve the outcomes for a variety of applications. As the low replicability or reliability is a common problem experienced by single-cell omics, our work may provide an effective tool to overcome this problem in single-cell omics analysis.

## Acknowledgements

This work is supported in part by the National Natural Science Foundation of China (61503314, 81788101, 81630042), Natural Science Foundation of Fujian Province (2019J01041), National Key R&D Program of China (2020YFA0803500), 111 Project (B12001), Research Unit fund from Chinese Academy of Medical Sciences (2019RU054).

## Author contributions

F.Z. and J.H. conceived the concept and supervised the project. F.Z. designed the model. F.Z. and X.K. wrote the code. F.Z., X.K. performed the experiments. F.Z., X.K., F.Y., T.C., and J.H. analyzed the experimental results. F.Z and J.H. drafted the manuscript.

## Methods

### Deep auto-regressive factor modeling

In the following, we provide a thorough description of the model design and architecture of scPheno^XMBD^. We assume that the majority of variations among cell can be attributed to the variability of a few factors (Fig. 1). In other words, each factor can induce heterogeneity among cells. The cell-to-cell variability is resulted from the combinatorial effect of these factors. Therefore, scPheno^XMBD^ describes the variations among cells by factor modeling, which allows for controlling the generation of gene expression with a few variables. We consider cell type, cell phenotype, and data batch as the factors responsible for the cell-to-cell variability. Let *y, k*_1_, and *k*_2_ denote cell type, cell phenotype, and data batch for a given cell *x*. scPheno^XMBD^ generate the gene expression profile of the cell *x* as follows (Supplementary Fig. 1):

a. Given the cell type annotation *y*, scPheno^XMBD^ infers the standard cell state for *y*, which can be considered as the archetypic cell for the given cell type.
b. scPheno^XMBD^ computes the difference between the cell-type annotation and the cell-type prediction *ŷ* for the cell *x*, which indicates the deviation *d*_*y*_ of the cell *x* from the archetypic cell in the latent state space. The label prediction is calculated via the Bayesian estimation given in the following section.
c. The actual state *z*_*y*_ of the cell *x* in the latent state space of the cell-type factor can be inferred from the cell-type annotation *y* and the deviation *d*_*y*_.

scPheno^XMBD^ repeats these steps to infer the actual states 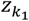 and 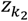 of the cell *x* in the latent state spaces corresponding to the cell-phenotype factor and the data-batch factor, respectively. The probabilistic description of the entire generative process is addressed in the follows,

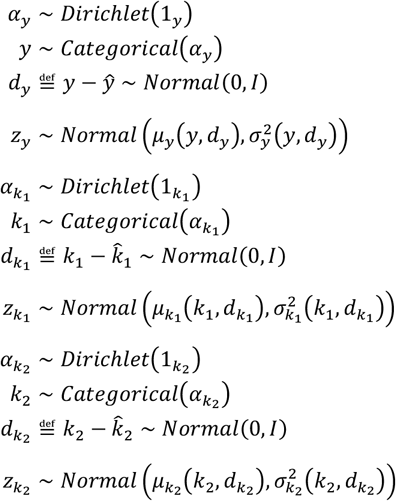

where, 1_*y*_, 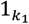, and 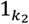 are the vectors of ones representing the non-informative prior probabilities over the parameters *α*_*y*_, 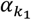, and 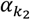 of the Categorical distributions. The mean and covariance vectors of the above equations are inferred by using multiple-layer neural networks. Another latent state variable *z*_*n*_ is added to account for the randomness in data. Therefore, the complete latent sate of the cell *x* is defined as,

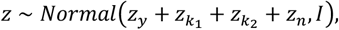

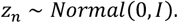

The expression of each gene *g* in the cell *x* is reconstructed by using zero-inflated negative binomial model (ZINB),

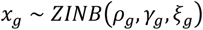

where *ρ*_*g*_ is the normalized gene frequency, *γ*_*g*_ is the gene-specific inverse-dispersion parameter, and *ξ*_*g*_ is the probability of the gene having zero expression due to dropout effects. The parameters *ρ*_*g*_ and *ξ*_*g*_ are inferred from the complete cell state *z* by using two multiple-layer neural networks *ρ* = P(*z*) and *ξ* = Ξ(*z*), whereas *γ*_*g*_ is the global variable in our implementation.

### Variational inference

scPheno^XMBD^ models the generation of single-cell gene expression with the auto-regressive factor model that involves the feedbacks of data annotation predictions. However, it is analytically intractable to compute the evidence *p*(*x*) for the calculation of the posterior distribution 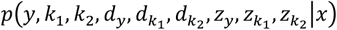 by using the Bayes’ rule. Instead, we utilize variational inference^28^ to approximate the posterior distribution with a variational distribution that can be factorized as the following,

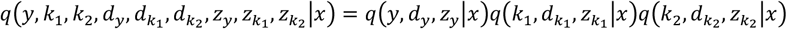

The variational distributions of distinct factors are independent to each other. The posterior distributions of the variables related to a specific factor can be approximated regardless of the other factors. For instance, the approximation of *p*(*y, d*_*y*_, *z*_*y*_|*x*) can be proceeded,

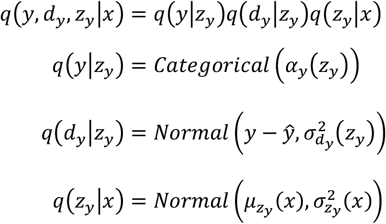

where the distribution parameters *α*_*y*_(*z*_*y*_), 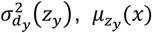, and 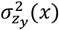 are estimated with multiple-layer neural networks. The variational distributions *q*(*y*|*z*_*y*_) and *q*(*z*_*y*_|*x*) can be used to predict the cell-type label *ŷ*) for the cell *x*,

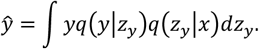

The posterior distributions 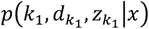 and 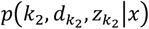 are approximated with the same procedure.

Both factor and variational models utilizes a number of multiple-layer neural networks to infer the distributions, which contain a considerable quantity of parameters to be optimized. To estimate the parameters of the factor model and the variational model, we use the ADAM optimizer^29^, a stochastic backpropagation algorithm to maximize the following evidence lower bound,

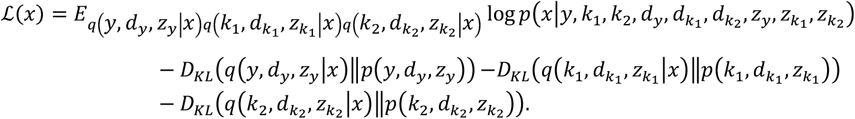

The model is implemented using Pytorch^30^. Our experiments are run on an NVIDIA TITAN GPU.

### Processing and analysis of the IFN*β* stimulation dataset

We utilized scRNA-seq data from ref.^19^ to illustrate the concept of gene expression factorization and extraction. The dataset was retrieved from the R package SeuratData under “ifnb”. The annotations of cell type, cell subtype, and stimulation status for each cell were made available by the data provider. We followed the standard instructions of Seurat to normalize and scale data, and then do PCA and UMAP analyses. Differential expression for scRNA-seq was conducted by using the FindAllMarkers function of Seurat with default parameters. The enrichment analysis of signaling proteins was performed by using the EnrichR^31^ website. Silhouette score is to measure the closeness of a cell to other cells with the same type and the separation of the cell to the other types. We used the Silhouette_samples function of the python package Scikit-learn^32^ to compute the Silhouette scores for cells. We used Silhouette scores to evaluate the changes in cell population structure.

### Processing and analysis of the human fibroblast dataset

We utilized scRNA-seq data of human fibroblast from ref.^23^ to illustrate the difficulty of identifying the set of genes that can consistently cluster cells. The data was normalized and processed following the standard instructions of Seurat. Different numbers of HVGs were generated by using Seurat’s function FindVariableFeatures with a variety of values for nfeatures. The clustering of cells was accomplished by using Seurat’s function FindClusters (resolution=1.0). The consistence between two clustering results was evaluated by using the python package Scikit-learn’s function adjusted_rand_score. Rand index is usually used to evaluate the information shared in two clustering results, but it prefers to assign a high score when both of two results contain numerous clusters. Adjusted Rand index accounts for the impact of the number of clusters, resulting in a more trustworthy evaluation.

### Processing and analysis of the Seok et al. dataset

We utilized scRNA-seq data of healthy participants and severe influenza patients from ref.^24^ to illustrate that the inter-subject discrepancy could hinder the interpretability of disease-association cell variations. The cell-type annotation and pathological status were made available by the data provider. To clarify the difficulty, CD14+ monocyte cells were extracted and processed using the standard Seurat protocol for displaying the combinatorial effect of inter-subject discrepancy and disease on cell states. We used Seurat’s function IntegrateData to remove the effect of inter-subject variations with the CCA algorithm. The integrated data was processed under the standard protocol of Seurat to demonstrate the result of data correction.

### Processing and analysis of the Ahern et al. dataset

We utilized scRNA-seq data of healthy participants and severe influenza patients from ref.^26^ to validate the generalizability of scPheno^XMBD^. The cell-type annotation and pathological status were made available by the data provider.

### Processing and analysis of the CRISPR screening dataset

We utilized scRNA-seq data from ref.^27^ to demonstrate the application of scPheno^XMBD^ on analyzing single-cell CRISPR screening data. The dataset was retrieved from the R package SeuratData under “thp1.eccite”. The annotations of edited gene and perturbation status for each cell were made available by the data provider. The data was processed according to the standard Seurat protocol.

## Data availability

All datasets used in the paper are public. Both IFN*β* stimulation and CRISPR screening datasets were retrieved from the R package SeuratData (“ifnb” and “thp1.eccite”). scRNA-seq data of human fibroblast cells were retrieved from Dr. Alkes Price’s laboratory website https://alkesgroup.broadinstitute.org/LDSCORE/Jagadeesh_Dey_sclinker. scRNA-seq of the Seok et al. dataset^24^ and the Ahern et al. dataset^26^ were retrieved from Human Cell Atlas Data Portal https://data.humancellatlas.org/explore/projects/95f07e6e-6a73-4e1b-a880-c83996b3aa5c and Zenodo https://doi.org/10.5281/zenodo.6120249, respectively.

## Code availability

scPheno^XMBD^ is available for academic use at https://github.com/ZengFLab/scPheno.

## Figure Legends

**Supplementary Figure 1.**
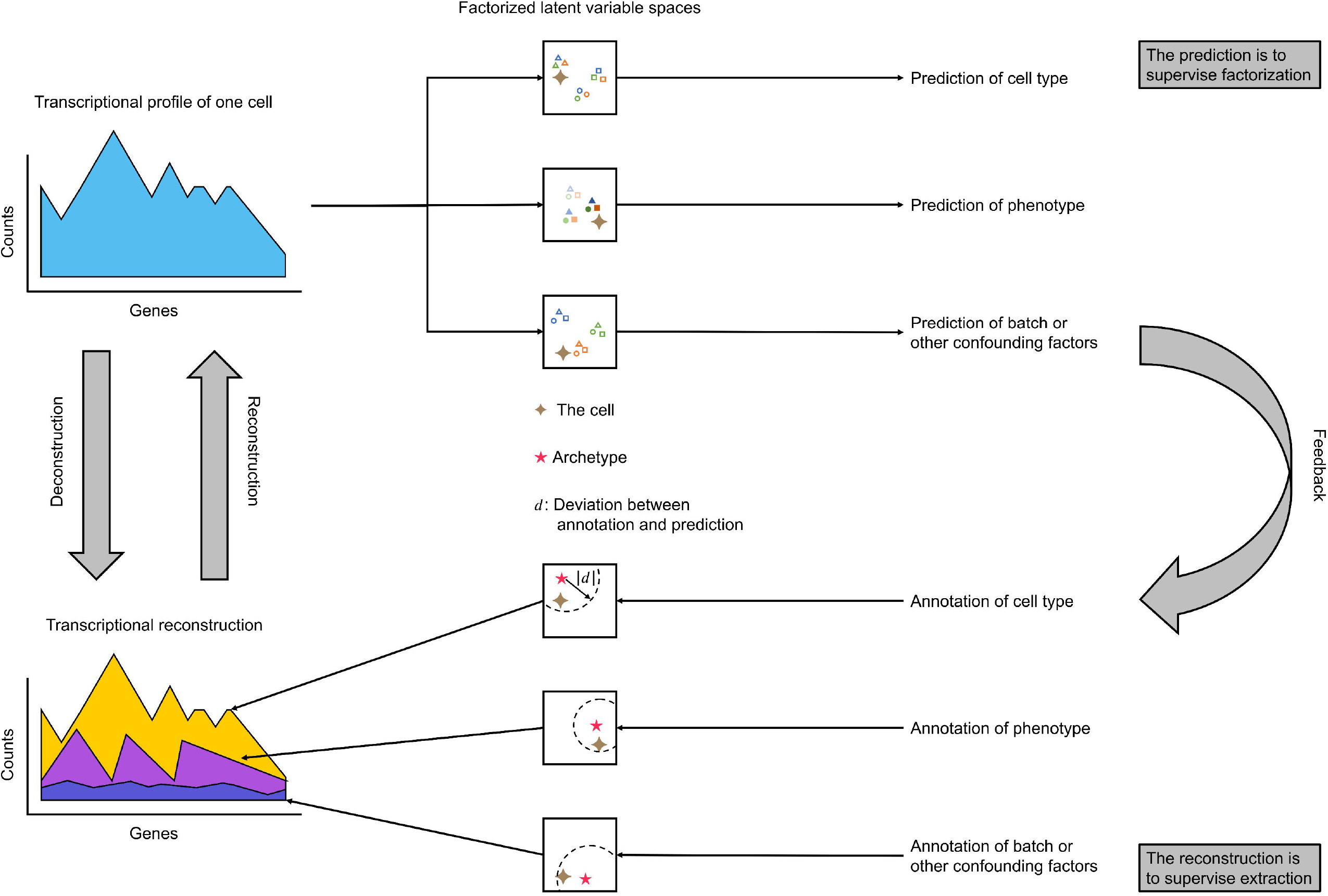
Schematic illustration of scPheno^XMBD^. scPheno^XMBD^ is a deep generative model based on factor modeling. scPheno^XMBD^ consists of the reconstruction and deconstruction components. **Top**, The deconstruction component disassembles variations in gene expression pertaining to factors, which then guide the reconstruction of cell state in the latent state space corresponding to each factor. **Bottom**, The reconstruction component infers the complete cell state by integrating cell states regarding various factors and then rebuild gene expression of the given cell based on the complete cell state.

**Supplementary Figure 2.**
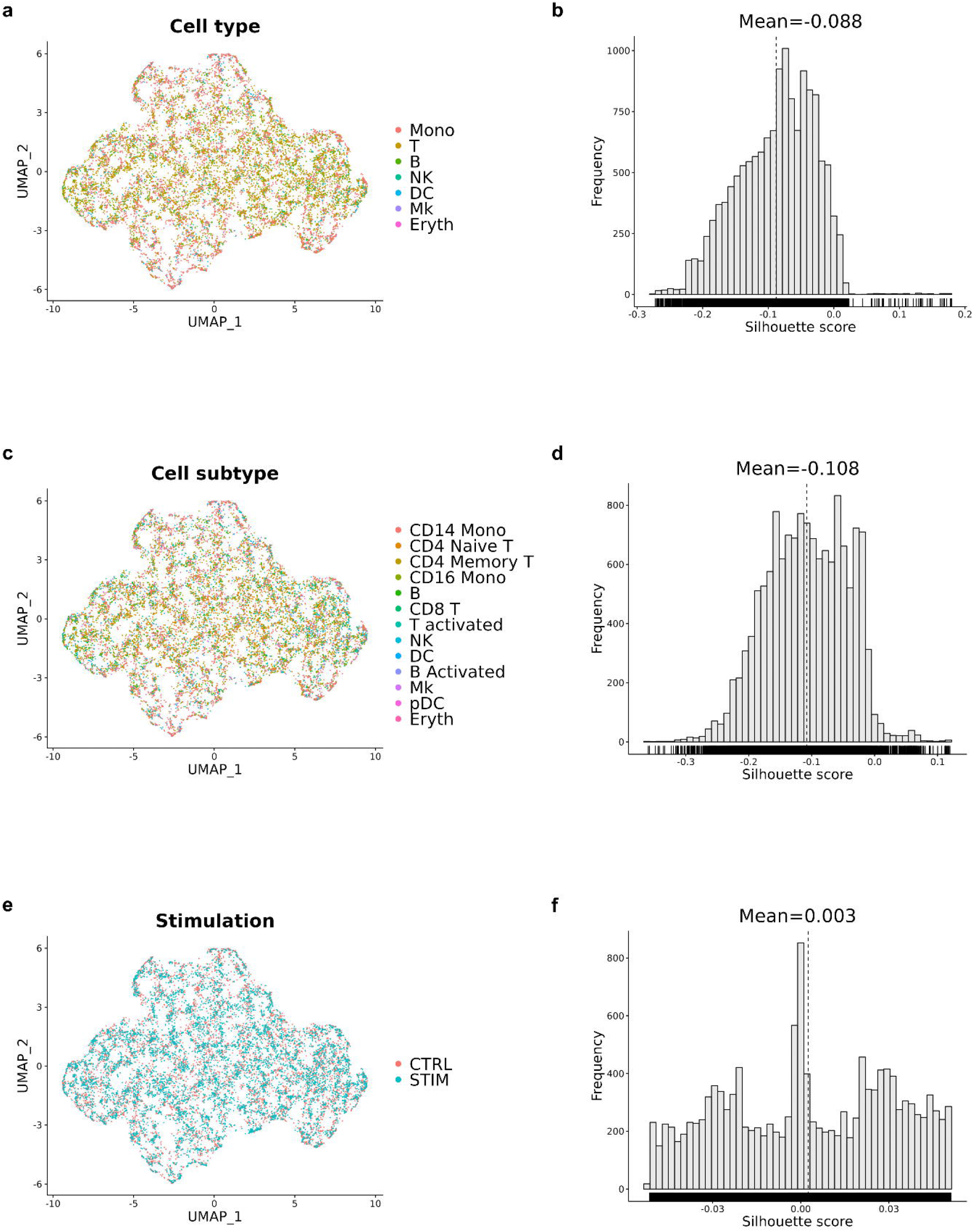
Cell distribution in the noise space. **a**,**b**, Cells are colored by cell type (**a**) and the separability of cell types evaluated by using the Silhouette scores (**b**). **c**,**d**, Cells are colored by cell subtype (**c**) and the separability of cell subtypes (**d**). **e**,**f**, Cells are colored by stimulation status (**e**) and the separability of stimulation status (**f**).

**Supplementary Figure 3.**
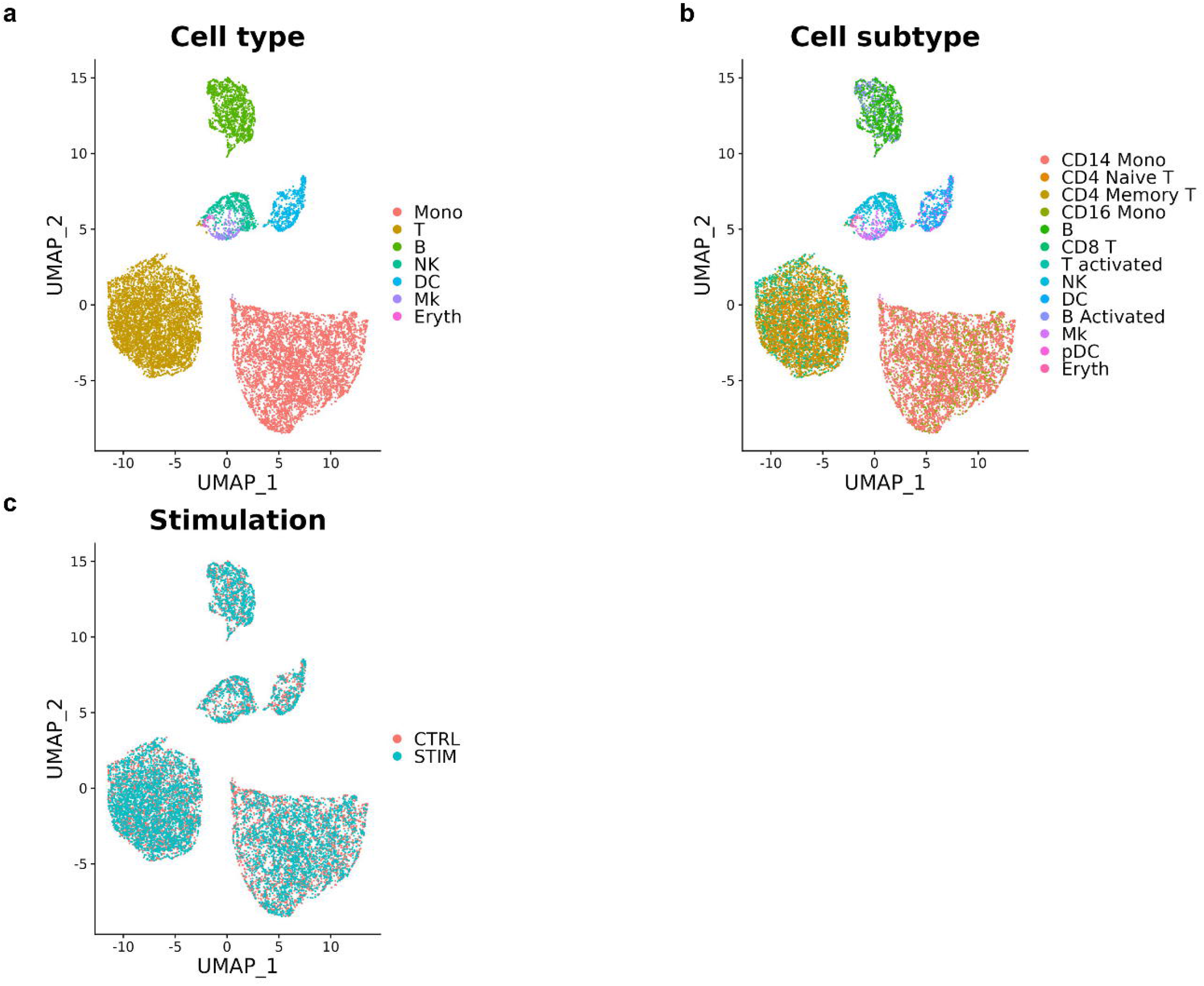
Cell distribution inferred from the reconstructed transcriptional profiles pertaining to cell type. **a-c**, Cells are colored by cell type (**a**), cell subtype (**b**), and stimulation status (**c**).

**Supplementary Figure 4.**
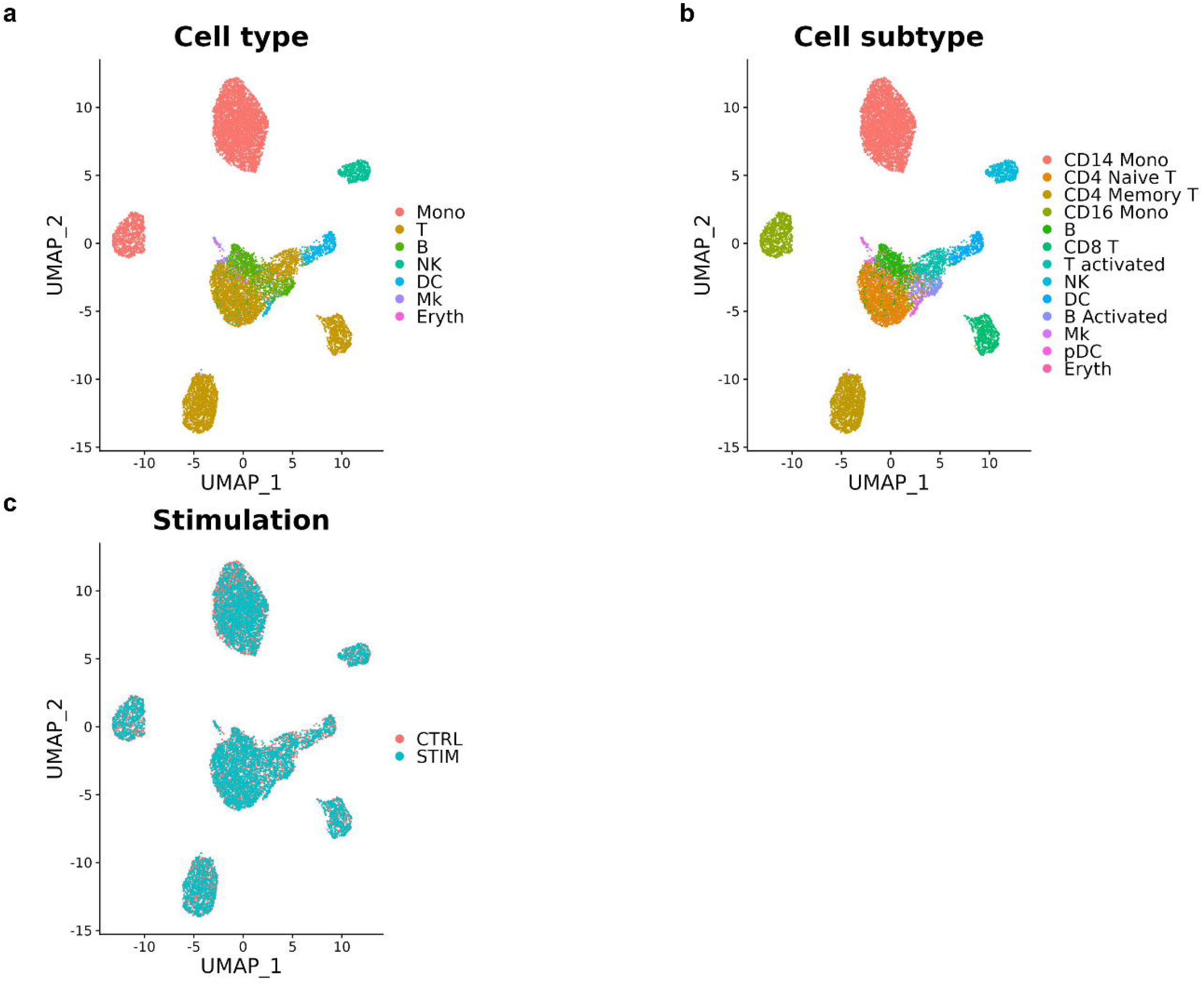
Cell distribution inferred from the reconstructed transcriptional profiles pertaining to cell subtype. **a-c**, Cells are colored by cell type (**a**), cell subtype (**b**), and stimulation status (**c**).

**Supplementary Figure 5.**
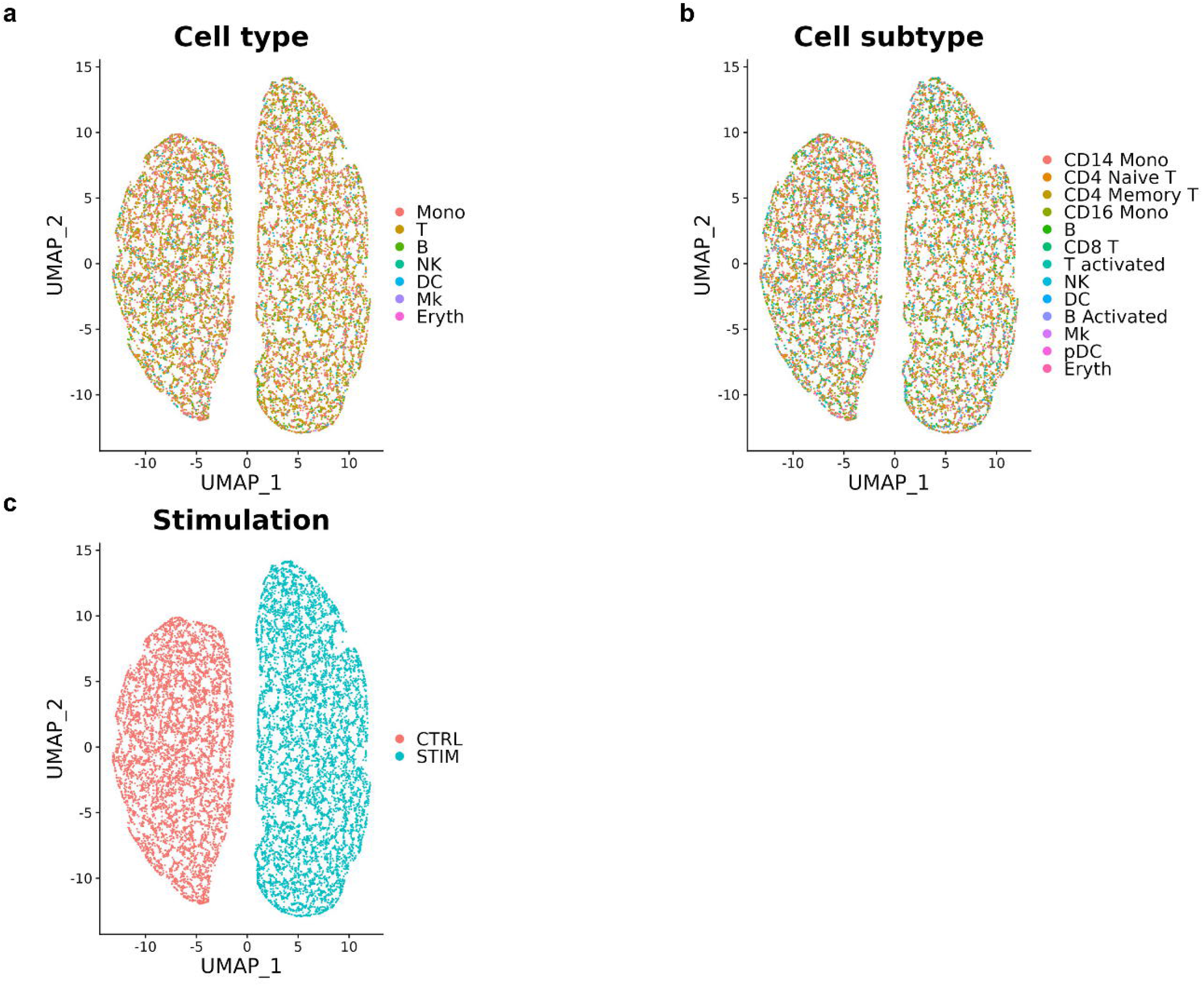
Cell distribution inferred from the reconstructed transcriptional profiles pertaining to stimulation status. **a-c**, Cells are colored by cell type (**a**), cell subtype (**b**), and stimulation status (**c**).

**Supplementary Figure 6.**
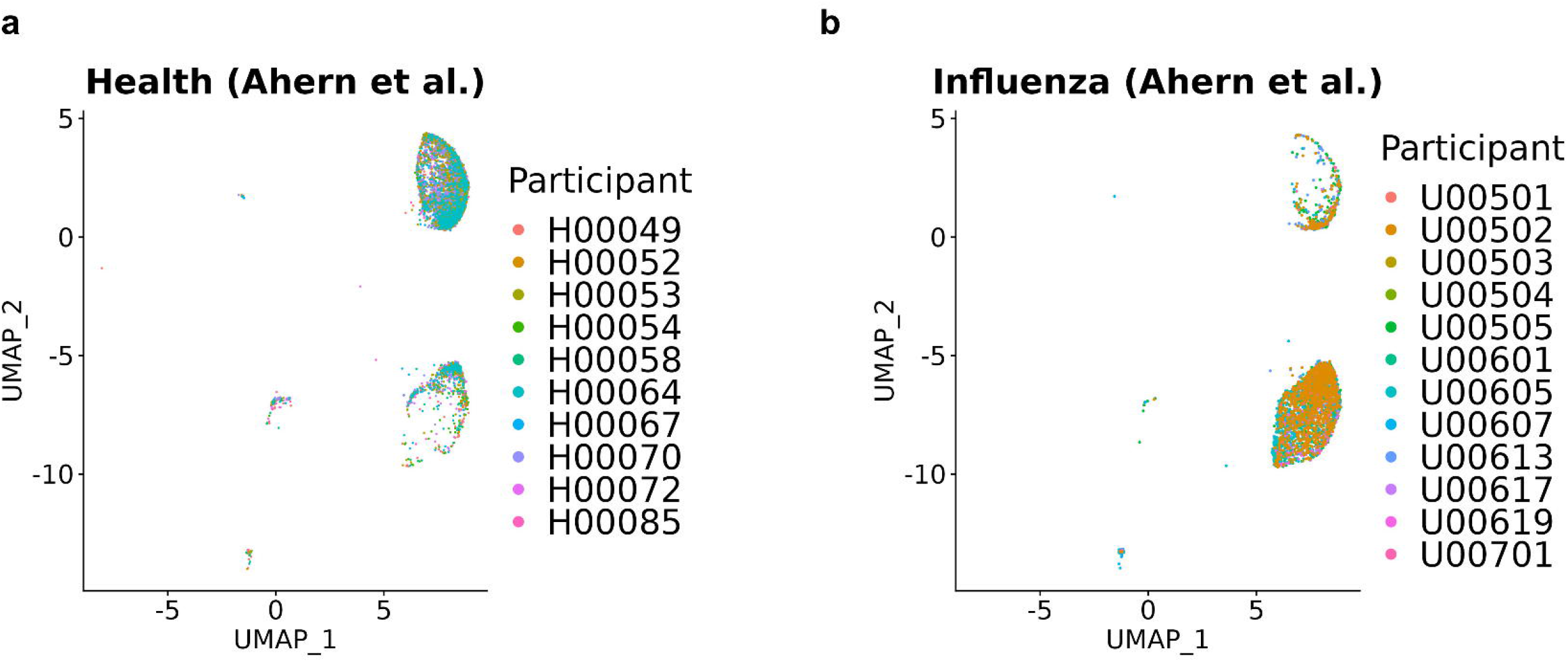
There are no inter-subject variations in the scPheno^XMBD^-processed the Ahern et al. dataset. **a**, The variations of the healthy participants among cells. **b**, The variations of the influenza infection patients among cells.

**Supplementary Figure 7.**
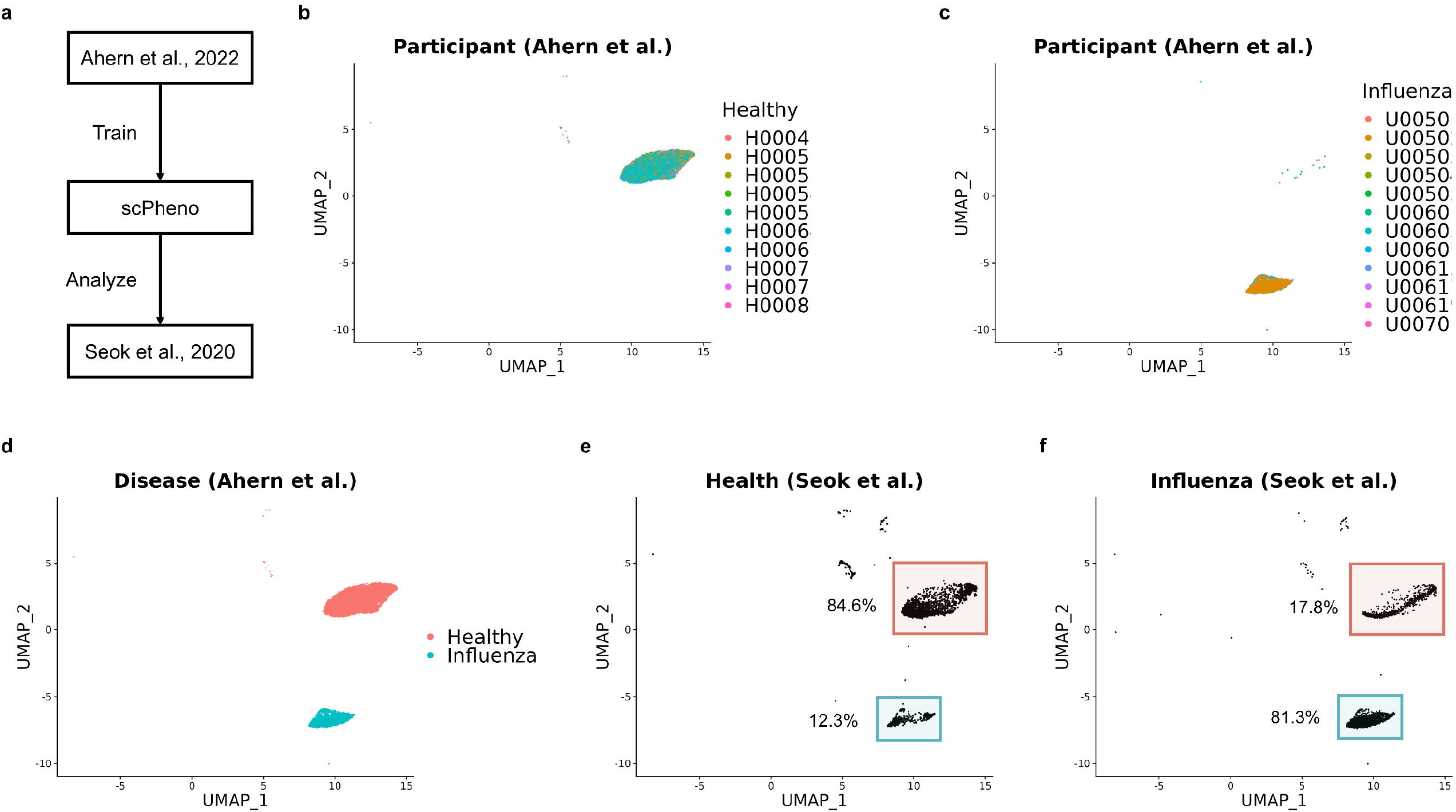
Validate the generalizability of scPheno^XMBD^. **a**, Apply the scPheno^XMBD^ model trained with the Ahern et al. dataset to the Seok et al. dataset. **b**,**c**, Eliminate the inter-subject variations of the Ahern et al. dataset in the healthy participants (**b**) and the influenza infection patients (**c**). **d**, scPheno^XMBD^ retains the variations between cells responsive to influenza infection. **e**, The embedding of CD14+ monocyte cells from the healthy participants in the Seok et al. dataset. The proportions of the correctly classified monocytes and misclassified CD14+ monocyte cells are approximately 84.6% and 12.3%, respectively. **f**, The embedding of CD14+ monocyte cells from the influenza infection patients in the Seok et al. dataset. The proportions of the correctly classified monocytes and misclassified CD14+ monocyte cells are approximately 81.3% and 17.8%, respectively.

